# Correcting batch effects in large-scale multiomic studies using a reference-material-based ratio method

**DOI:** 10.1101/2022.10.19.507549

**Authors:** Ying Yu, Naixin Zhang, Yuanbang Mai, Qiaochu Chen, Zehui Cao, Qingwang Chen, Yaqing Liu, Luyao Ren, Wanwan Hou, Jingcheng Yang, Huixiao Hong, Joshua Xu, Weida Tong, Leming Shi, Yuanting Zheng

## Abstract

Batch effects are notorious technical variations that are common in multiomic data and may result in misleading outcomes. With the era of big data, tackling batch effects in multiomic integration is urgently needed. As part of the Quartet Project for quality control and data integration of multiomic profiling, we comprehensively assess the performances of seven batch-effect correction algorithms (BECAs) for mitigating the negative impact of batch effects in multiomic datasets, including transcriptomics, proteomics, and metabolomics. Performances are evaluated based on accuracy of identifying differentially expressed features, robustness of predictive models, and the ability of accurately clustering cross-batch samples into their biological sample groups. Ratio-based method is more effective and widely applicable than others, especially in cases when batch effects are highly confounded with biological factors of interests. We further provide practical guidelines for the implementation of ratio-based method using universal reference materials profiled with study samples. Our findings show the promise for eliminating batch effects and enhancing data integration in increasingly large-scale, cross-batch multiomic studies.

---

Batch effects are notorious technical variations irrelevant to study factors of interests, but are common in transcriptomics^1-4^, proteomics^5, 6^, metabolomics^7^, and multiomics integration^8^. Due to variation in experimental design, lab conditions, reagent lots, operators, and other non-biological factors, results from different batches may vary and result in misleading outcomes^9-14^.

Batch effects can have a particularly negative impact on study outcomes^9, 10^. On the one hand, the presence of batch-correlated variation could skew analysis and introduce large numbers of false-positive and false-negative findings, and even misleading conclusions^15^. For example, a change of experiment solution caused a shift in the risk calculation, leading to incorrect treatment decision^16^. On the other hand, systematic variations including batch effects become one of the major causes of irreproducibility^17,18^. What is worse, reproducibitily crisis raises questions about the reliability of data. For example, some researchers repeat their microarray sample profiling with RNAseq to avoid batch effects introduced by the two platforms. Such a costly undertaking could be averted when data from distinct platforms can be integrated properly^19, 20^. With the era of big data, the issue of batch effects becomes more prominent^10, 21^.

Although many batch-effect correction algorithms (BECAs) have been proposed^7, 12, 13, 22-25^, studies that comprehensively assess the performances of various BECAs for application to multiomic studies are currently lacking, or is still controversial. For example, in transcriptomics, several widely used BECAs, such as ComBat^7, 22^, Surrogate variable analysis (SVA)^23^, and RUVseq^24^, have been shown acceptable performances in some researches^26-28^, but did not perform well in others^10, 29^. Similarly, ratio by calibration with common reference sample(s), which is also known as Ratio-G, has shown improved comparability in some multi-batch studies^2, 30, 31^, and less comparability in other studies^27^. Recently, Harmony, a principal component analysis (PCA) reduction-based method, was shown to perform well in batch-group balanced and confounded scenarios in single-cell RNAseq data^32, 33^. But whether Harmony works well in other types of data remains to be seen.

Furthermore, datasets used for comparisons in previous evaluation studies are insufficient to determine the nature of batch effects. Several studies were based on heterogeneous samples^33, 34^, which were difficult to assess pure batch variations against hidden subpopulation variabilities among batches. And some studies were based on simulated datasets^27, 34^, which do not necessarily accurately represent the true nature of batch effects. These datasets used in the previous studies could not objectively reflect the nature of batch effect and might bring biases the evaluation of the performances of BECAs. Therefore, studies based on real-world, cross-batch datasets are urgently needed for objective assessment of BECAs.

Moreover, the levels of confounding between biological and batch factors may greatly influence the validity of BECAs. In a balanced scenario where study samples are evenly distributed across batch factors, batch effects can be mitigated via diverse BECAs^10, 27, 35^. The balanced scenario is ideal but almost impossible in reality. In most cases, biological factors and batch factors are often mixed and difficult to distinguish, which is recognized as the confounded scenario commonly seen in longitudinal and multi-center cohort studies. When biological factors and batch factors are strongly confounded, most BECAs may no longer be applicable^10, 27^. Therefore, there is an urgent need to identify batch correction methods to facilitate the integration of datasets from confounded batch-group scenarios.

Here, as part of the Quartet Project for quality control and data integration of multiomic profiling, we comprehensively assess the performances of seven BECAs for mitigating the impact of batch effects in multiomic datasets, including transcriptomics, proteomics, and metabolomics data. We previously established the Quartet RNA^36^, protein^37^ and metabolite^38^ reference materials derived from the same B-lymphoblastoid cell lines from the four members of a monozygotic twin family^39^. Large number of multiomic datasets were generated from multiple labs, platforms, and protocols. These rich datasets provide a unique opportunity for us to objectively assess the performances of BECAs based on the nature of batch effects under both balanced and confounded scenarios. The performances are evaluated in terms of accuracy of identifying differentially expressed features, robustness of predictive models, and classification accuracy after multiomic data integration. Our findings show the promise for eliminating batch effects and enhancing data integration in increasingly large-scale, cross-batch multiomic studies.

## Results

### Overview of the study design

Advantages and limitations of BECAs under balanced and confounded scenarios were shown in **Fig. 1a**. Suppose we have a total of 12 samples from two groups (A and B), including six As and six Bs, and the objective is to detect differentially expressed features (DEFs) between group A and group B. Ideally, in a balanced scenario where the two batches contain an equal number of samples from both groups A and B, batch effects can be effectively removed by many batch-effect removal methods, such as mean-centering per feature per batch. However, experimental scenarios are rarely balanced. In an extreme scenario when sample group is completely confounded with batch number in that all six As are processed in one batch and all six Bs in another batch, it is almost impossible to distinguish the real biological differences between A and B from technical variations resulting from batch effects. In this case, an incorrect combination of scenario-methods can lead to false negatives, because the true biological differences between the two groups could be removed together with batch effects. An effective way of tackling batch effects is to profile the same reference materials (e.g., Quartet multiomic reference materials) along with study samples in each batch. Expression profiles of each sample can be transformed to ratio-based values using reference samples as the denominator, whether in balanced or confounded scenarios (**Fig. 1a**).

**Figure 1.**
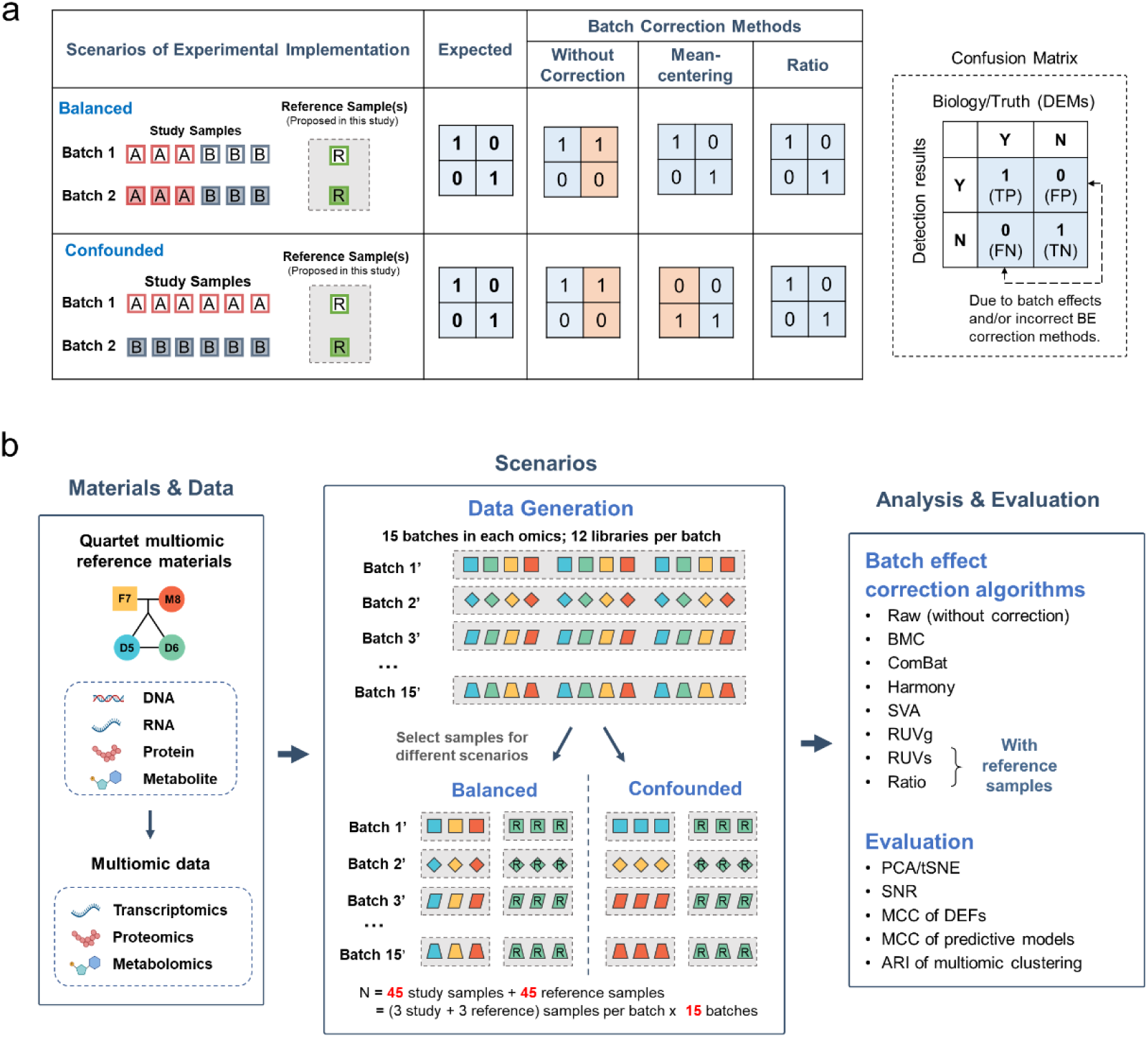
Overview of the study design. (**a**) Advantages and limitations of batch-effect correction algorithms (BECAs) under balanced and confounded experimental scenarios. False positives and false negatives in cross-batch comparisons using different BECAs. (**b**) Overview of datasets and analysis approaches. Multi-batch datasets from transcriptomics, proteomics, and metabolomics were generated using the Quartet multiomic reference materials derived from a Quartet family including Father (F7), Mother (M8), and monozygotic twin daughters (D5 and D6). Subsets of data were selected from the full datasets to create balanced and confounded scenarios for assessing the pros and cons of BECAs under these experimental scenarios. The multiomic profiling data were analyzed with seven BECA methods. Performances were then evaluated using visualization tools and quantitative metrics.

To objectively assess performances of BECAs, multiomic and multi-batch datasets based on the Quartet reference materials were used (**Fig. 1b**). As described in accompanying papers^36-38, 40, 41^, a set of DNA, RNA, protein, and metabolite reference materials was established simultaneously from four immortalized B-lymphoblastoid cell lines (LCLs) derived from a Quartet family including monozygotic twin daughters (D5 and D6) and their father (F7) and mother (M8). Reference materials were then distributed to multiple labs for generating multiomic data. In each omic type, 12 samples in 12 tubes with each representing one of the triplicates of reference sample groups were used for data generation in a batch concurrently. On the other hand, high-throughput experiments at different time points, in different labs, using different platforms or experimental protocols are recognized broadly as cross-batch experiments. Finally, multiomic datasets, including transcriptomics, proteomics, and metabolomics datasets from multiple labs, platforms, protocols, and batches were obtained, comprising a total of 252 RNA libraries from 21 batches^36^, 312 protein libraries from 26 batches^37^, and 204 metabolite libraries from 17 batches^38^.

We then selected a subset of datasets from the full datasets to create balanced and confounded scenarios for assessing the pros and cons of BECAs (**Fig. 1b**). In the balanced experiment scenario, 15 batches of data were used for each omic type, with each batch containing three study groups (D5, F7 and M8) and one replicate per group. In the confounded experiment scenario, the same 15 batches of data were used for each omic type, with each batch containing three replicates from one of the three study groups (D5, F7, or M8). Additionally, three replicates from the reference group (D6) in each batch were employed for reference-based BECAs. Therefore, 45 study samples and 45 reference samples in balanced and confounded scenarios were employed at each omic level (**Fig. 1b** and **Supplementary Table 1)**.

We evaluated seven BECAs, including mean-centering (BMC), ComBat^22^, Harmony^42^, SVA^23^, RUVg^24^, RUVs^24^ and ratio-based scaling (see Methods for details). We visualized clustering projections with both PCA and t-distributed stochastic neighbor embedding (t-SNE). We also applied four quantitative metrics for performance evaluation. First, signal-to-noise ratio (SNR) is used for quantifying the ability to separate biological groups when multiple batches of data are integrated. Secondly, correlation coefficient between a dataset and the reference datasets in terms of log2 fold changes. Thirdly, Matthews’ correlation coefficient (MCC) of a dataset using the reference dataset of differentially expressed feature (DEFs) as the truth. Reference datasets were generated from the consensus of differentially expressed features from intra-batch profiling. Fourthly, MCC was used to represent the predictivity of models for predicting the sex and age of the donors from whom the reference materials were derived. Finally, adjusted rand index (ARI) is used for measuring the accuracy of classification after multiomic data integration (**Fig. 1b**).

### Multiomic measurements are prone to batch effects and can be corrected using appropriate methods

We first applied PCA scatter plots to visualize the biological and (or) batch effects. In transcriptomics, it could be observed that, without correction, experimental factors rather than biological groups (D5, F7, or M8), exhibited the largest differences. BMC and ComBat performed well in distinguishing samples according to their biological groups only in the balanced scenario, not in the confounded scenario. In contrast, four BECAs, including two BECAs with reference samples (RUVs and ratio-based scaling), RUVg and SVA performed equally well in both balanced and confounded scenarios (**Fig. 2a** and **Extended Data Fig. 1a**). Similar results were observed in proteomics (**Fig. 2b** and **Extended Data Fig. 1b**). In metabolomics, Harmony, SVA, RUVg and RUVs did not perform as well as in transcriptomics, probably because they were developed primarily with transcriptomic data (**Fig. 2c** and **Extended Data Fig. 1c**). The straightforward methods such as BMC and ratio-based scaling were omics-independent, which means that similar performances were observed in transcriptomics, proteomics, and metabolomics.

**Figure 2.**
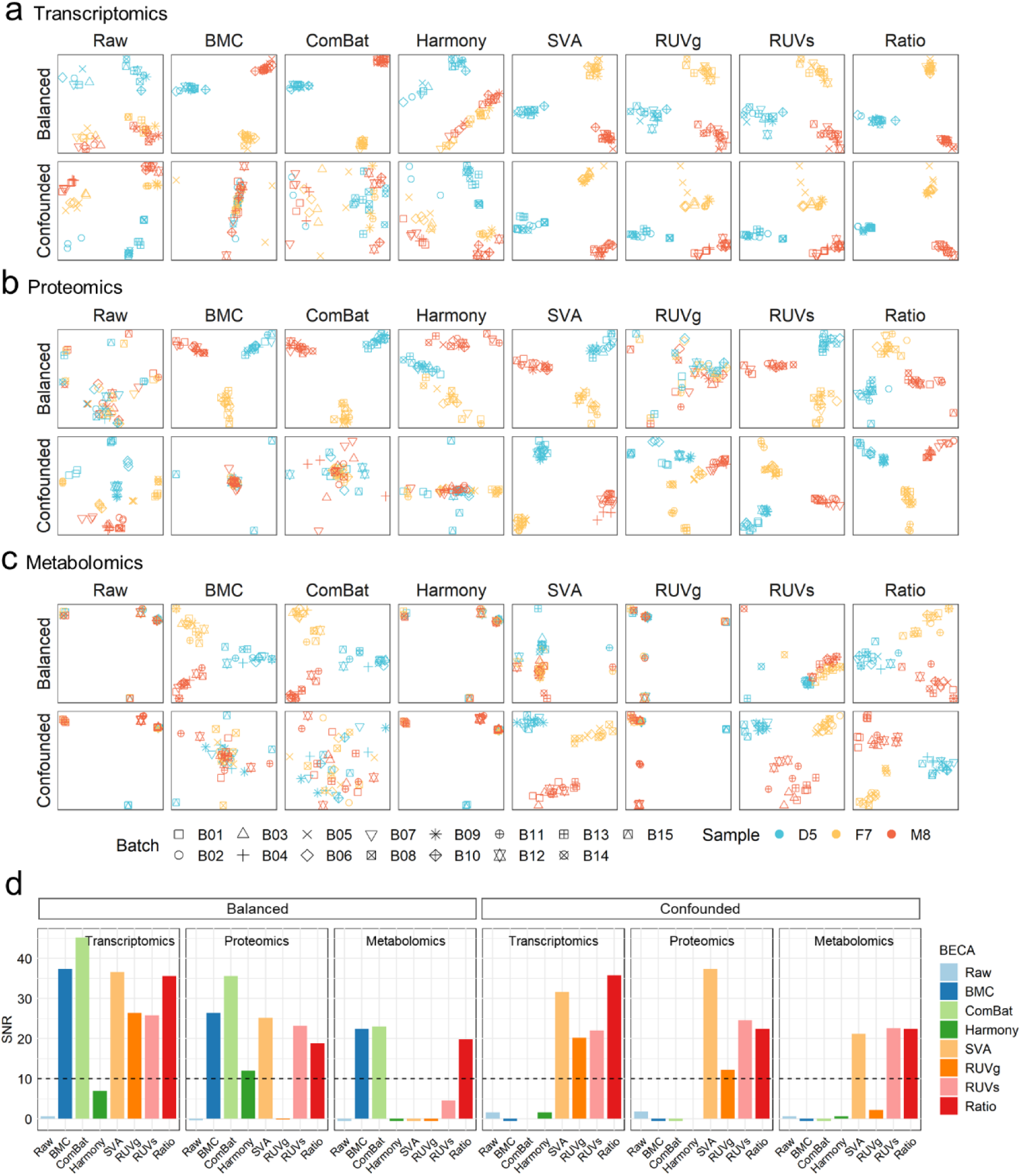
Multiomic measurements are prone to batch effects and can be corrected using appropriate methods. (**a-c**) PCA plots based on different batch-effect correction algorithms in balanced and confounded scenarios, using transcriptomics (**a**), proteomics (**b**), and metabolomics (**c**) data. Plots were color-coded by sample group (D5, F7, and M8), and shaped by batch. (**d**) Bar plot of signal-to-noise ratio (SNR) using different batch-effect correction methods on transcriptomics, proteomics, and metabolomics data.

PCA based performance metric Signal-to-Noise Ratio (SNR) was then used to quantify differences between biological sample groups and variations in technical replicates. SNR measures the ability of distinguishing intrinsic biological differences among distinct sample groups (“signal”) from technical variations including batch effects of the same sample group (“noise”), as mentioned in the accompanying papers^36, 39^. Generally, a higher SNR value indicates higher discriminating power, vice versa. SNR values were consistently high in ratio-based scaling in balanced and confounded scenarios among the three types of omics, whereas SNR values of SVA, RUVg and RUVs were high for one or two types of omics, but low for the others. On the other hand, SNR values of BMC and ComBat were high for balanced scenario but low in confounded scenario in all three types of omics (**Fig. 2d**).

### Accuracy of identifying differentially expressed features

As identifying differentially expressed features (DEFs) is one of the most important tasks for quantitative omics, we compared DEF identification performances across batches, using the consensus of intra-batch DEFs and the average of fold-changes between sample-pairs as “ground truth” for benchmarking. We then developed two quality metrics, relative correlation and MCC of DEFs. Specifically, we introduced the “Relative correlation” (RC) metric, i.e., the Pearson correlation coefficient between the fold-changes of a test dataset for a given pair of samples and the corresponding intra-batch fold-changes, and “MCC of DEFs” (MCC) metric, i.e., Matthews Correlation Coefficient (MCC) to measure the consistency of DEFs detected from a test dataset for a given pair of samples with those from the common intra-batch DEFs (**Fig. 3a**).

**Figure 3.**
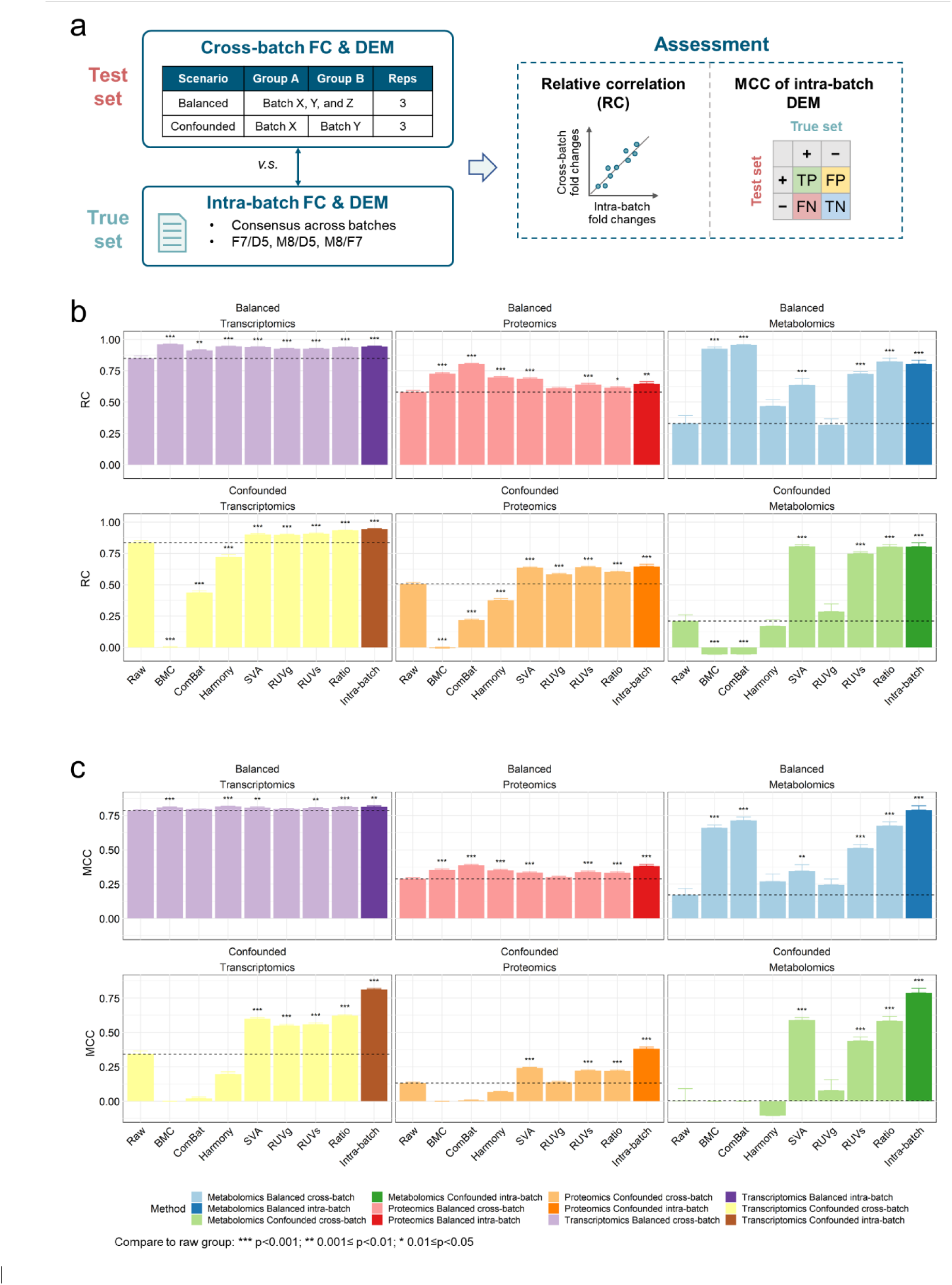
Accuracy of differentially expressed features. (**a**) Schematic diagram of comparisons of differentially expressed features (DEFs) between cross-batch tests and reference datasets. (**b-c**) Bar plots (mean ± s.e.) representing the relative correlation (RC) (**b**) and Matthews Correlation Coefficient (MCC) of DEFs (**c**) with reference datasets and across seven BECAs in balanced and confounded scenarios using transcriptomics, proteomics, and metabolomics data. Mean of the raw data group in each panel was plotted in dashed line. T-test between each BECA method with the raw data group was conducted. Symbolic number coding of *p*-value was used as: *** (*p*≤0.001), ** (*p*≤0.01), * (*p*≤0.05).

Using RC values as a metric, BMC, ComBat, SVA and ratio-based scaling gave significantly higher RC values compared to cross-batch DEFs without correction across three omics in balanced scenario. On the contrary, SVA, RUVs and ratio-based scaling gave significantly higher RC values in confounded scenario (*p*<0.05) (**Fig. 3b**). Particularly, in metabolomic data in confounded scenario, RC increased from 0.13 ± 0.27 to 0.81 ± 0.10 (mean ± SD) after conducting ratio-based scaling. Moreover, some BECAs were able to improve RC to values as high as intra-batch RC, while others significantly reduced RC values, highlighting the importance of choosing a suitable BECA, especially in confounded scenario (**Fig. 3b**). Using MCC of DEFs as a metric, SVA and ratio-based scaling consistently outperformed other methods, which was in line with the assessment by RC (**Fig. 3c**). Indeed, when applying a variety of widely used performance metrics, such as sensitivity, specificity, precision, and Jaccard Index of DEFs, we observed similar performances across different BECA methods (**Extended Data Fig. 2**), indicating the difficulty of using these metrics to differentiate performances of various BECAs.

### Accuracy of model prediction

Cross-batch prediction is another important task in quantitative omics, especially in the context of biomarker discovery for clinical diagnosis, prognosis, and therapeutic action. Thus, we evaluated the impact of BECAs on cross-batch prediction.

Frequently, a predictive model was built using a set of samples, and was further validated using independent datasets[cite MAQC-II main paper and Luo J batch-effect paper]. These datasets could be confounded with batch effects. In this study, we divided samples into two sets before developing predictive models. Specifically, 27 libraries from nine batches were used as training set and 18 libraries from six batches as validation set, according to data generation date, as we did in a previous publication^4^. The training set was then used to select variables and train prediction models using five machine-learning algorithms, including model averaged neural network (avNNet), support vector machine (SVM), random forest (RF), generalized partial least squares (GPLS), and linear algorithm BstLm, through an internal-layer of 25 runs of fivefold cross-validation process to resist overfitting. The model was further validated using the validation set as an external-layer evaluation. Age and sex of the donors from whom the Quartet reference materials were developed were used as biological endpoints to assess the robustness of cross-batch prediction. These two endpoints (age and sex) are easy to predict compared to most clinically relevant endpoints. Thus, the failure of accurate prediction of these easy endpoints implies serious problems in clinical settings.

Based on multiple evaluation metrics, we found that modeling performances for the balanced scenario were equally good, regardless of the choice of BECA methods (**Fig. 4** and **Extended Data Fig 3**). On the other hand, for the confounded scenario, SVA, RUVg, RUVs and ratio-based scaling performed well, whereas BMC, ComBat and Harmony performed as bad as or worse than non-correction (**Fig. 4** and **Extended Data Fig 3**). This trend remained consistent for transcriptomics, proteomics, and metabolomics.

**Figure 4.**
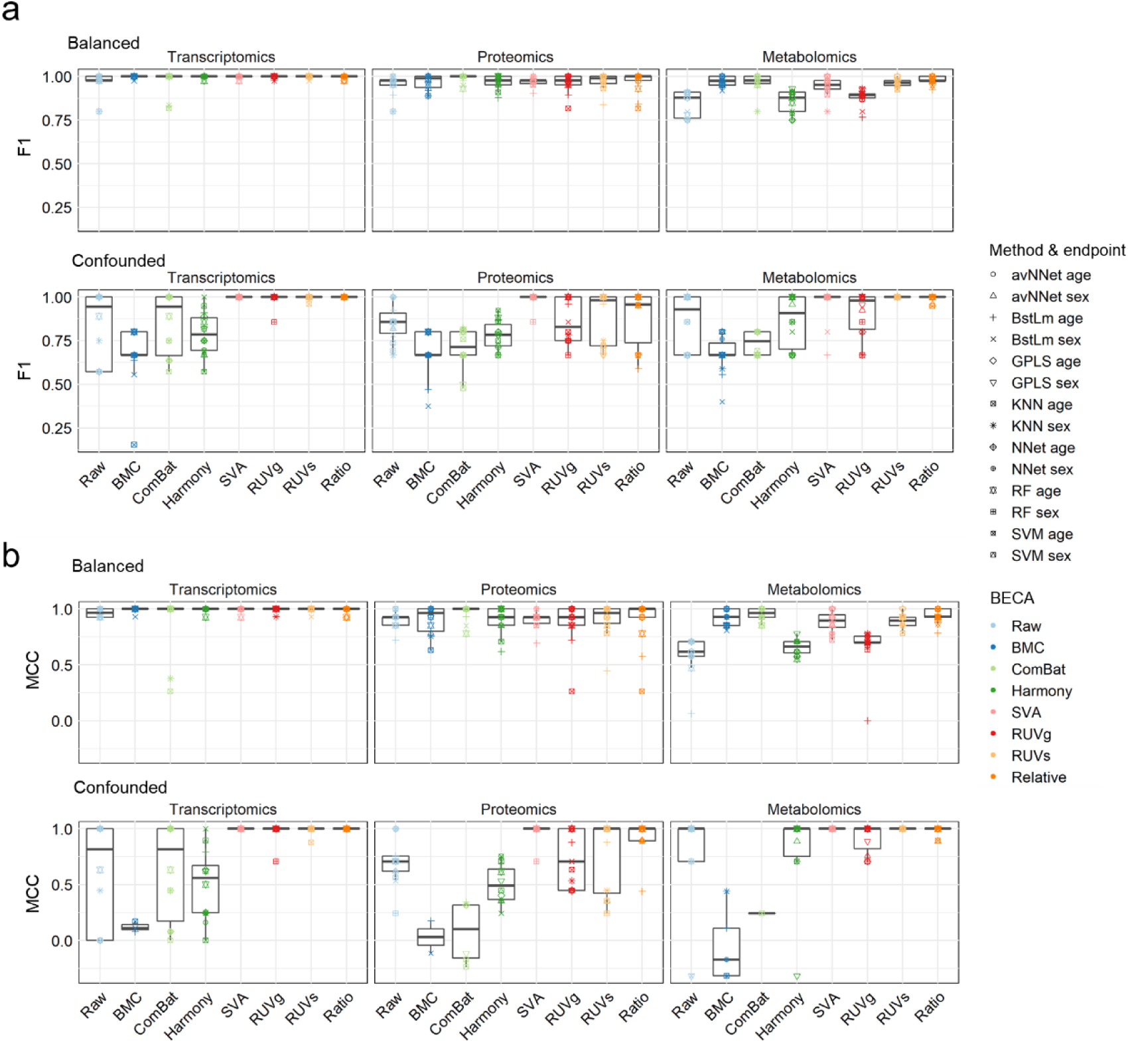
Accuracy of model prediction. Boxplots of modeling performances in predicting sex and age using five machine-learning algorithms under bot balanced and confounded scenarios. Performances were measured using F1 scores (**a**) and MCC (**b**).

### Consistency of multiomic clustering

As clustering multiomic data has the potential to find disease subtypes and to reveal systems level insights, it has become one of the most popular applications in integrative analysis. Hence, we further compared performances of these BECAs in terms of ability of accurately clustering cross-batch samples into their biological sample groups (D5, F7 and M8) after multiomic data integration. Datasets consisting 36 samples of three sample groups derived from 12 batches were randomly selected from the entire dataset (see Methods for details). Three widely-used integrative tools were used, including SNF^43^ (**Fig. 5a**), intNMF^44^ (**Fig. 5b**), and iClusterBayes^45^ (**Fig. 5c**). The performance was measured using the Adjusted Rand Index (ARI)^46^, a commonly used metric to compare the clustering labeling against the true labeling.

**Figure 5.**
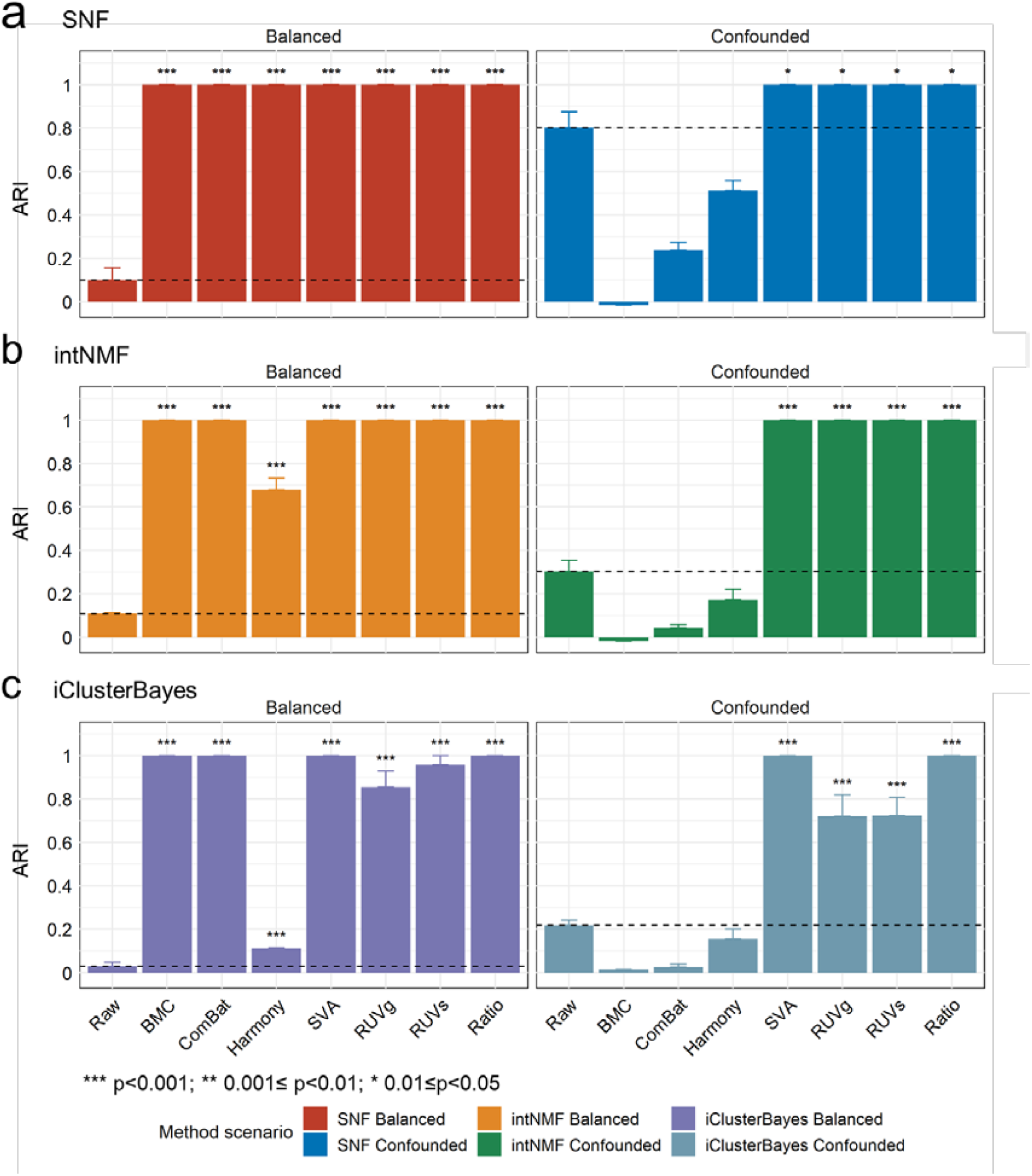
Consistency of multiomic clustering according to their sample groups. Bar plots (mean ± s.e.) of Adjusted Rand Index (ARI) values of multiomic clustering using different batch-effect correction algorithms in balanced and confounded scenarios. Three integrative tools were used, including SNF (**a**), intNMF (**b**), and iClusterBayes (**c**). Expression profiles from 36 samples from three sample groups derived from 12 batches in each omic type were randomly selected from the entire datasets and further used to integrate cross-omics data. In order to eliminate random effect, the random selection and cross-omics integration was conducted for ten times. Average value of the raw data group in each panel was plotted in dashed line. T-test between each BECA method with the raw data group was conducted. Symbolic number coding of *p*-value was used as: *** (*p*≤0.001), ** (*p*≤0.01), * (*p*≤0.05).

SVA and ratio-based scaling consistently outperformed other BECAs across three omics types (**Figs. 5a-c**). Additionally, RUVg and RUVs performed well in transcriptomics and proteomics, but were less effective in metabolomics. BMC and ComBat showed excellent performance (ARI = 1) in the balanced scenario; however, they performed poorly (ARI around zero) in the confounded scenario. The choice of different integrative tools made little differences to the results except for datasets after Harmony correction. Our results highlighted the problems of widely used BECAs in real-world scenarios where batch effects are prevalent.

### Overall performances of BECAs

We provided a summary of the overall performances of seven BECAs as measured by SNR, MCC for DEFs, MCC for model prediction, and ARI for multiomic data integration (**Fig. 6a**). Ratio-based scaling ranked on the top and revealed a general superiority by significant improvements in SNR, identification of DEFs, prediction and multiomic clustering, compared to raw data without correction. Besides, SVA, RUVs, and RUVg were alternative methods that were suitable in both balanced and confounded scenarios. ComBat and BMC were highly context-dependent and were only suitable in the balanced scenario. Harmony, a BECA method developed based on single-cell RNAseq data, showed limited improvement in bulk RNAseq, proteomics, and metabolomics data.

**Figure 6.**
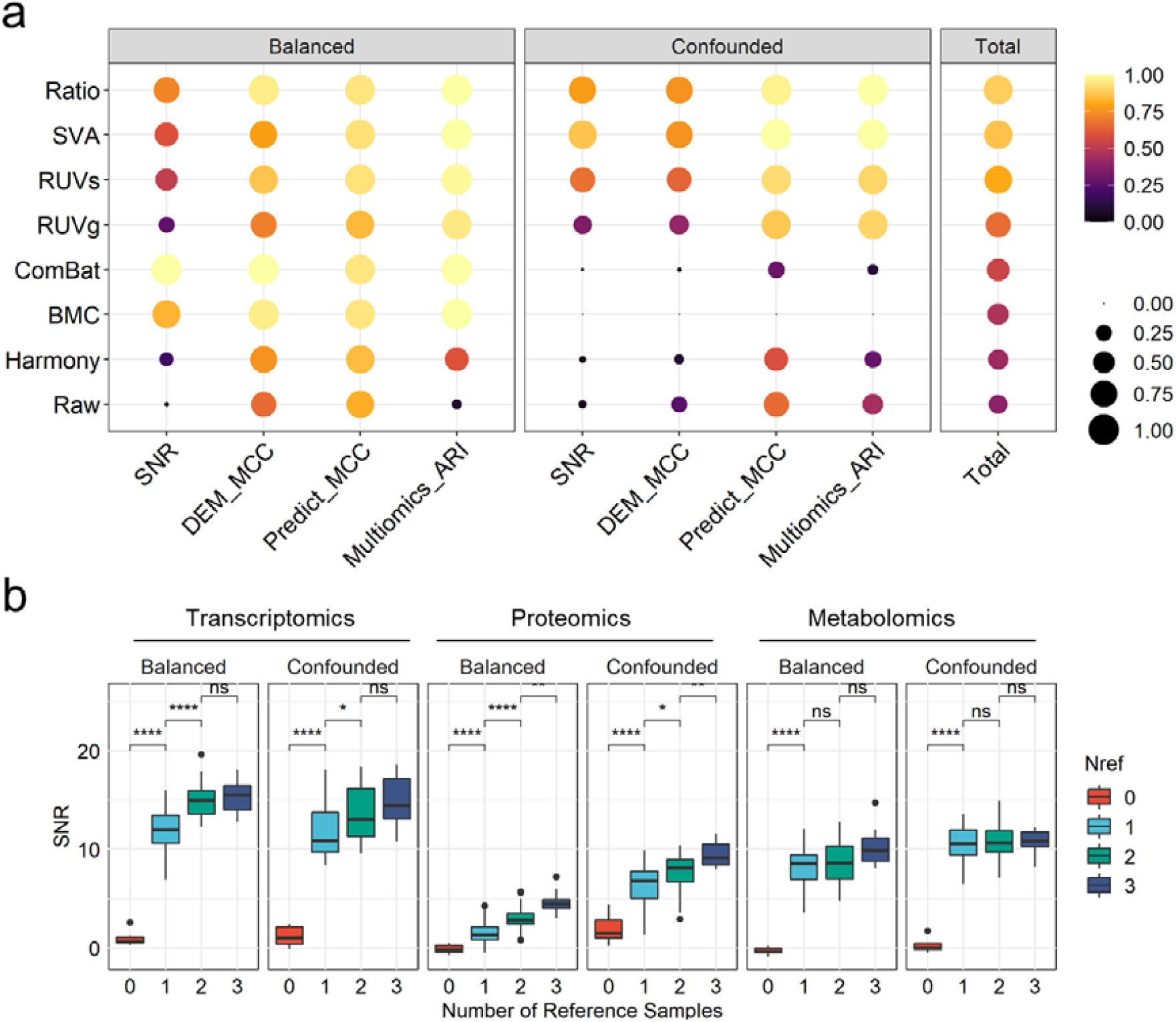
Summary of performances of BECAs and choice of number of samples for ratio-based scaling. (**a**) The summarized performance of seven BECAs in balanced and confounded scenarios. The BCCAs were ordered by their total score. For the calculation of the total score, we first separately scaled the values of the four metrics, including Signal-To-Noise Ratio (SNR), Matthews Correlation Coefficient (MCC) of identification of differentially expressed features (DEFs), MCC of prediction, and Adjusted Rand Index (ARI) of multiomic clustering to an interval of (0,1) to equalize the weight of different metrics. The total score was expressed as mean of the scaled values of the four metrics. (**b**) Boxplot of Signal-to-Noise Ratio (SNR) under different numbers of replicates of the reference sample used as dominators in conducting ratio-based scaling. T-test was conducted. Symbolic number coding of *p*-value was used as: *** (*p*≤0.001), ** (*p*≤0.01), * (*p*≤0.05), ns (*p*>0.05, not significant).

### The choice of number of samples for ratio-based scaling

If we used reference materials to conduct ratio-based scaling, an important question is what number of samples would constitute an appropriate choice as the denominator for converting absolute expression data to ratio-based scales. Thus, the number of reference samples that could be used as a reference in ratio-based scaling within each batch were tested. As expected, SNR increased when using ratio-based expressions compared to absolute expressions even when only one sample with one replicate was used, and further increased when more replicates were added to calculate the average expression values as the denominator (**Fig. 6b**). These findings emphasized that it is critical to use reference samples per-batch along with study samples, and that it is better to use more reference samples.

## Discussion

Batch effects in multiomic profiling are universal and detrimental to study purpose. Our results showed that batch effects were prevalent in quantitative profiling technologies, presenting challenges for combining data from different batches of single-omics and multiomics. Hence, batch correction is an essential step in multi-batch analysis.

Applying BECAs is highly context-dependent. In a balanced scenario, the batch effects are evenly distributed across study groups and can be got rid of via all seven BECAs we tested. In reality, the ideal batch-group design is almost impossible in multi-center and longitudinal cohort studies, when batch effects can be fully confounded with the investigational endpoints of interests. Furthermore, batch effects hamper the legitimacy of retrospective data integration aiming to explore new insights from comparison of several independent cohort studies, such as the healthy and disease cohorts^10^. In these cases, some BECAs, such as BMC and ComBat, were no longer applicable. What is worse, incorrect usage of BECAs could lead to many adverse effects such as removal of true biological signals.

Our results indicated that the application of ratio-based method was warranted. We prefer the ratio-based method for three reasons. First, the ratio-based method is easy to implement, platform-independent, and applicable to multiomics quantification, including transcriptomics, proteomics, and metabolomics. Secondly, compared to ComBat or BMC, the ratio-based scaling is less affected by study design of unbalanced distributions of samples in different groups between different batches. In clinical applications and large-scale projects, the imbalance of samples across different batches is inevitable. Thirdly, compared to the ratio-based scaling, SVA may remove potentially important biological information encoded in the latent variables.

According to our results, using two or three replicates of a common reference material in each batch and converting expression data to feature-wise ratio-based scaling profiles within each batch can play an important role in making expression levels more comparable and resistant to batch effects. As the Quartet multiomic reference materials and the corresponding reference datasets have been successfully developed in our accompanying work^36-39^, which represent the first suites of publicly available multiomic reference materials, we therefore recommend the use of the Quartet reference materials for monitoring and correcting batch effects.

Our study bears some limitations. First, the number of samples used for developing predictive models was small and the biological endpoints (sex and age) were relatively easy, which could not fully represent clinical applications. However, the performances presented here could be considered as an upper bound of the respective methods being investigated. Secondly, samples used in the study were derived from the Quartet reference materials. Although clear trend of pros and cons across BECAs could be observed, larger sample size and more tissue types of samples should be further investigated.

In summary, multiomic measurements are prone to batch effects, which, fortunately, can be effectively monitored and corrected by using ratio-based scaling of the multiomic data. Profiling common reference materials concurrently with study samples can enhance data comparability of multi-batch studies, especially for large-scale multiomic studies.

## Methods and Materials

### Multiomic datasets from the Quartet reference materials

To examine impact of batch factors in multiomic data, we used reference materials and datasets from the Quartet Project. Establishment of Epstein-Barr virus (EBV) transformed B-lymphoblastoid cell lines and multiomic reference materials, as well as implementation of high-throughput experiments were described in an accompanying paper by Zheng et al^39^. Briefly, multiomic reference materials (DNA, RNA, protein, and metabolite) were derived from immortalized Epstein-Barr Virus (EBV) infected B-lymphoblastoid cell lines from a four-member Chinese Quartet family including two monozygotic twin daughters (D5 and D6), and their father (F7) and mother (M8) with expected differences in multiomic profiles between any two members^36-38, 40^.

Reference materials were then distributed to multiple labs for generating multiomic profiling data. According to the Quartet Project study design, in each omics, 12 samples were used as a standard sample set, consisting of 12 tubes with each representing one of the triplicates of a reference sample group^39^. The high-throughput experiments were conducted concurrently for the 12 samples. On the other hand, high-throughput experiments at different time points, in different labs, using different platforms or experimental protocols are recognized broadly as cross-batch experiments. The high-throughput datasets, metadata, and primary analyses are deposited in the Quartet Data Portal (http://chinese-quartet.org/) and described in an accompanying paper by Yang et al.^41^.

### Data generation, analysis, and normalization

Data generation, analysis, and normalization of transcriptomics, proteomics, and metabolomics data were described in accompanying papers^36-41^.

#### Transcriptomics

A total of 252 whole transcriptome sequencing (RNAseq) libraries from eight labs with either poly-A selection or rRNA-removal protocol were constructed. On average, 100 million read-pairs per replicate were sequenced on Illumina NovaSeq or MGISEQ-2000. Detailed information was described in the accompanying RNA paper ^36^.

RNAseq reads were aligned using HISAT2 and genes were quantified using StringTie followed by Ballgown^47^. The Fragments Per Kilobase of transcript per Million mapped reads (FPKM) is used to represent the normalized abundance of each gene. A gene was considered detectable if the normalized FPKM value was higher than 0.1 in over 30% of the samples and was retained for further analysis. A floor value of 0.01 was added to the FPKM value of each gene before log2 transformation.

#### Proteomics

A total of 312 LC-MS/MS datasets were generated under a data-dependent acquisition mode (DDA) using different platforms. The MS platforms included Thermo Fisher Scientific™ Q Exactive™ hybrid quadrupole-Orbitrap™ series mass spectrometers (Q Exactive, Q Exactive Plus, Q Exactive HF and Q Exactive HF-X), Thermo Fisher Scientific™ Orbitrap Fusion™ Tribrid™ series mass spectrometers (Fusion and Fusion Lumos), Orbitrap Exploris 480 mass spectrometer, Sciex Triple-TOF 6600 and Bruker timsTOF Pro mass spectrometer. Detailed information was described in the accompanying protein paper^48^.

MS raw files were searched against the National Center for Biotechnology Information’s (NCBI) human Refseq protein database (updated on 04-07-2013, 32,015 entries) using Firmiana 1.0 enabled with Mascot 2.3 (Matrix Science Inc)^49^. False discovery rate (FDR) by using a target-decoy strategy was set to 1% for both proteins and peptides. Proteins were then quantified using the label-free iBAQ approach. The fraction-of-total (FOT) is used to represent the normalized abundance of a particular protein, which was defined as a protein’s iBAQ value divided by the total iBAQ of all identified proteins within one sample^49^. A protein was considered detectable if the normalized FOT value was higher than 0.1 in over 30% of samples and retained for further analysis. A floor value of 0.01 was added to the FOT value of each protein before log2 transformation.

#### Metabolomics

The dried cell extracts were re-dissolved in mobile phase in each lab, and a total of 204 libraries were generated from five labs. The non-targeted metabolomics datasets were generated using AB SCIEX Triple TOF 5600, AB SCIEX QTRAP 6500, AB SCIEX TripleTOF 6600, and Thermo Fisher Scientific Q Exactive HF hybrid quadrupole-Orbitrap mass spectrometer systems in four difference labs. The targeted metabolomics datasets were generated using Waters Xevo TQ-S and AB SCIEX QTRAP 6500 mass spectrometers in two labs. More information was detailed in the accompanying metabolite paper^50^.

A metabolite was considered detectable if the normalized value was higher than 1 in over 70% of samples and retained for further analysis. A floor value of 1 was added to the value of each metabolite before log2 transformation.

### Datasets in balanced and confounded scenarios

In this study, we selected 45 samples from 15 batches as study samples, including three samples in each batch from three sample groups (D5, F7, and M8). In each batch, three samples from one replicate of each sample group (D5-1, F7-1, and M8-1) were used in the balanced scenario, whereas three samples from three replicates of one sample group (D5/F7/M8) were used in the confounded scenario. Additionally, 45 samples of D6 from 15 batches (three replicates of D6 in each batch) were used as reference samples in both balanced and confounded scenarios (**Supplementary Table 1**). This experimental design ensured the consistent number of samples in the balanced and confounded scenarios, as well as the separation of study samples from the reference samples, ensuring objective evaluation of the impact of batch effects and BECAs.

### Batch-effect correction methods

#### Batch Mean-centering (BMC)

Mean-centering per feature per batch is to set the mean of each feature across all the samples within each batch to zero. This approach is applied in all transcriptomics, proteomics, and metabolomics datasets based on log2 transformed expressions.

#### ComBat/ComBat-seq analysis

ComBat is one of the most popular BECA tools^7^. It applies empirical Bayes shrinkage to adjust the mean and the variance by pooling information across multiple genes for correcting batch-effects^22^. ComBat-seq extends ComBat adjustment framework to using negative binomial regression to estimate RNAseq count data^7^. The ComBat function in the sva 3.42.0 package^7^ was implemented for normalized expressions of proteomics and metabolomics, while the ComBat_seq function in the ComBat-Seq package^23^ was implemented for count data of transcriptomics.

#### Harmony analysis

Harmony uses an iterative clustering-correction procedure based on soft clustering to correct for sample differences. The algorithm first combines the batches and projects the data into a dimensionally reduced space using PCA, and then uses an iterative procedure to remove the batch effects. The HarmonyMatrix function in the harmony 0.1.0 package^42^ was implemented, using default parameter settings.

#### Surrogate variable analysis (SVA)

SVA is able to remove unwanted sources of variation while protecting the contrasts due to the primary variables specified in the function call. The sva functions in the sva v3.42.0 package^23^ was implemented to detect and remove latent variables, using default parameter settings.

#### Remove unwanted variation (RUV)

Remove unwanted variation (RUV) uses a subset of the data to estimate the factors of unwanted variation adjusts for nuisance technical effects. We applied two modes for estimating the factors of unwanted variation, including: (1) RUVg, using negative control genes, assumed not to be differentially expressed with respect to the covariates of interest; and (2) RUVs, using reference samples (D6) for which the covariates of interest are constant^24^. The RUVg and RUVs functions in the RUVSeq v1.28.0 package^24^ was implemented, using default parameter settings.

#### Ratio-based scaling

Ratio-based scaling refers to converting expression profiles to relative-scale profiles within each batch on a feature-by-feature basis. Ratio-based scaling data were obtained by subtracting log_2_-transformed expression profiles of a feature by the mean of log_2_-transformed expression profiles of the three replicates of D6 in the same batch.

### SNR

Signal-to-noise ratio (SNR) is defined as the ratio of the average distance among different samples (like D5-1 vs F7-1) from the average distance among technical replicates of the same sample (like D5-1 vs D5-2). Based on principal component analysis (PCA), distances of two samples in the space defined by the first two PCs were used to represent distances between the two samples. SNR was calculated as follows:

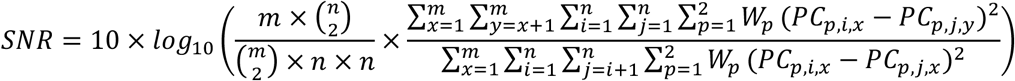

where *m* is the number of sample groups, and *n* is the number of replicates in each sample group. *W*_*p*_ represents the p^th^ principal component of variances. *PC*_*p,i,x*_, *PC*_*p,j,x*_ and *PC*_*p,j,y*_ represent the p^th^ component values of replicate *i* and replicate *j* in sample group *x* or sample group *y*, respectively.

### Intra-batch fold changes

Reference fold-changes at intra-batch level were defined from consensus integration of multi-batch datasets from transcriptomics, proteomics, and metabolomics, providing “ground truth” for benchmarking. Briefly, we defined the intra-batch fold-changes of sample-pairs (F7/D5, M8/D5, and M8/F7) in the format of mean by summarizing from 21, 26 and 17 batches of transcriptomics, proteomics, metabolomics data respectively for generating high-confidence intra-batch fold changes. In order to improve the reliability of the fold-changes, features that were satisfied with thresholds of detectability and t-test *p* < 0.05 in over 15% of the batches across the two samples in each sample.

### Intra-batch differentially expressed features (DEFs)

A list of high-confidence DEFs and non-DEFs were identified from consensus integration of multi-batch datasets. A gene/protein/metabolite was considered as a high-confidence DEF between two sample types if it was concordantly discovered as up- or down-regulated (*p* < 0.05 and fold change ≥ 2 or ≤ 0.5) in more than 15% of batches. Additionally, a gene/protein/metabolite was considered as a high-confidence non-DEF between two sample types if it was not differentially expressed (t-test *p* > 0.05 and fold change < 2 and > 0.5) in more than 50% of batches.

### Relative correlation

Relative correlation was calculated based on the Pearson correlation coefficient between the ratio-based expression levels of a dataset for a given pair of groups and the corresponding intra-batch fold-change values. It is referred to as the “relative correlation” metric, representing the numerical consistency of the ratio-based expression profiles. To improve reliability, the mean of the three replicates of each sample group was calculated before performing ratio-based expression analysis. Fold-changes were transformed using log2 scaling.

### MCC of DEFs

Cross-batch DEFs were calculated based on normalized expression profiles. Features with t-test *p* < 0.05 and fold change ≥ 2 or ≤ 0.5 were labeled as up-regulated or down-regulated, respectively. Based on high-confidence DEFs and non-DEFs, we can calculate the number of true positive (TP), true negative (TN), false positive (FP), and false negative (FN), and calculate the MCC to measure the consistency of DEFs detected from a dataset for a given pair of samples with those from the high-confidence DEFs. This metric is called the “MCC of DEFs”.

### Prediction models

Frequently, a predictive model was built using a set of samples, and was further validated using independent datasets. These datasets were probably confounded with batch effects. To simulate the clinical context, age and sex corresponding to the sample of each library were used as biological endpoints to assess the accuracy of cross-batch prediction.

Prediction models were developed and validated using a two-layer validation strategy^4^. Briefly, samples were first divided into two sets, comprising 27 libraries from nine batches as training set, and the remaining 18 libraries from six batches as validation set, according to data generation date. The training set was then used to select variables and train prediction models using five machine-learning algorithms, including model averaged neural network (avNNet), support vector machine (SVM), random forest (RF), generalized partial least squares (GPLS), and linear algorithm BstLm, through an internal-layer of 25 runs of fivefold cross-validation process to resist overfitting. The model was further validated using the validation set as an external-layer evaluation. Model performance was assessed using multiple performance metrics, including F1 score, MCC, sensitivity, specificity, PPV (positive prediction value), NPV (negative prediction value), precision, and accuracy. Model construction and assessment was implement using caret package v6.0-90 (https://github.com/topepo/caret).

### Integration of multiomic data

Expression profiles from 36 samples of three sample groups derived from 12 batches in each omics type were randomly selected from the entire dataset (N=45) and further used to integrate cross-omics data. In order to eliminate random effect, this process was repeated ten times. Three integrative tools were used, including iClusterBayes from iClusterPlus v1.30.0 package^51^, intNMF from IntNMF 1.2.0 package^52^, and SNF from SNFtool 2.3.1 package^43^. The number of eigen features of iClusterBayes was set to 3. The number of clusters of IntNMF was set to 3. Parameters for SNF were set as follows: the number of neighbors package (12), hyperparameter (0.5), the number of iterations (10), and the number of clusters (3). All other parameters were set by default.

The Adjusted Rand Index (ARI) was used to measure consistency of clustering after multiomics integration with true sample labeling. The Rand Index (RI) computes a similarity measure between clusters by considering all pairs of samples and counting pairs that are assigned in the same or different clusters in the predicted and true clusters. The raw RI score is then “adjusted for chance” into the ARI score as follows:

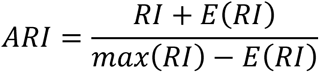

### Quantification and statistical analysis

All statistical analyses and data visualization were implemented using R statistical packages (4.1.2) (https://www.r-project.org). Student’s t-test was used to compare continuous variables. PCA was conducted with the univariance scaling, using the *prcomp* function. tSNE was conducted using the R package Rtsne v0.15. Data visualization was implemented using the R packages ggplot2 v3.3.5 (https://ggplot2.tidyverse.org/), GGally v2.1.2 (http://ggobi.github.io/ggally/), and ggsci v2.9 (https://github.com/nanxstats/ggsci). JMP^®^ Pro v16.0.0 (SAS Institute Inc., Cary, NC, USA) was used for predictive modeling analysis.

## Disclaimer

The content is solely the responsibility of the authors and does not necessarily represent the official views of the US Food and Drug Administration.

## Acknowledgments

This study was supported in part by National Key R&D Project of China (2018YFE0201603 and 2018YFE0201600), the National Natural Science Foundation of China (31720103909 and 32170657), Shanghai Municipal Science and Technology Major Project (2017SHZDZX01), State Key Laboratory of Genetic Engineering (SKLGE-2117), and the 111 Project (B13016). Some of the illustrations in this paper were created with BioRender.com.

## Author contributions

Y.Z., L.S., W.T, and Y.Y. conceived and oversaw the study. Y.Y., N.Z., Y.M., Q.C., Z.C., Q.W.C., Y.L., L.R., W.H., J.Y., H.H., J.X., W.T., L.S., and Y.T.Z performed data analysis and/or data interpretation. Y.Y., N.Z, Y.M., Y.Z., L.S., W.T., J.X., and H.H. wrote and/or revised the manuscript. All authors reviewed and approved the manuscript. Dozens of participants of the Quartet Project freely donated their time and reagents for the completion and analysis of the project.

## Competing interest declaration

The authors declare no competing financial interests.

## Additional information

### Extended Data Figures and Legends

**Extended Data Fig. 1.**
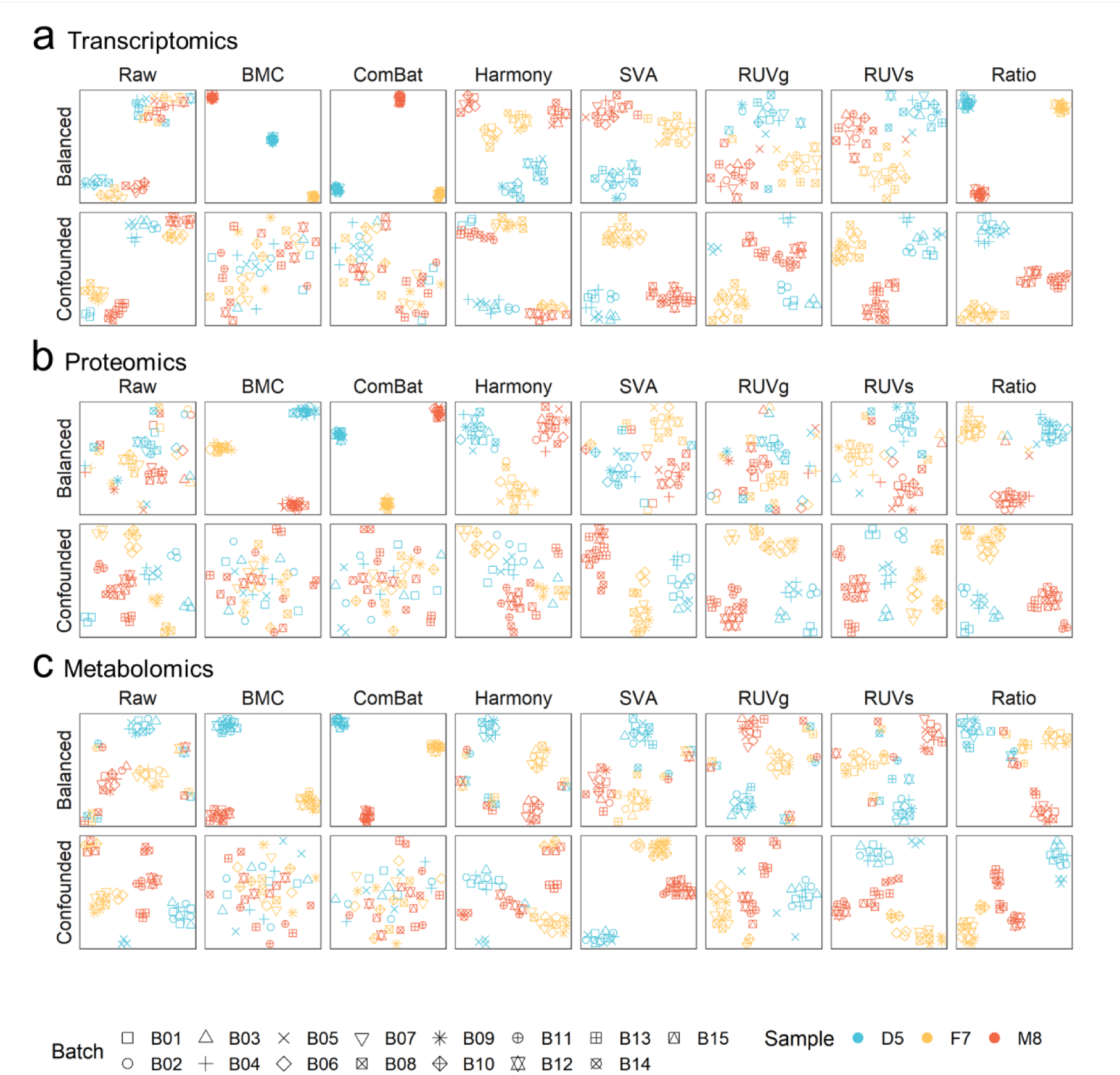
tSNE plots based on different batch-effect correction methods in balanced and confounded scenarios. Transcriptomics (**a**), proteomics (**b**), and metabolomics (**c**) data were used. Plots were color-coded by sample group (D5, D6, F7, and M8) and shaped by batch.

**Extended Data Fig. 2.**
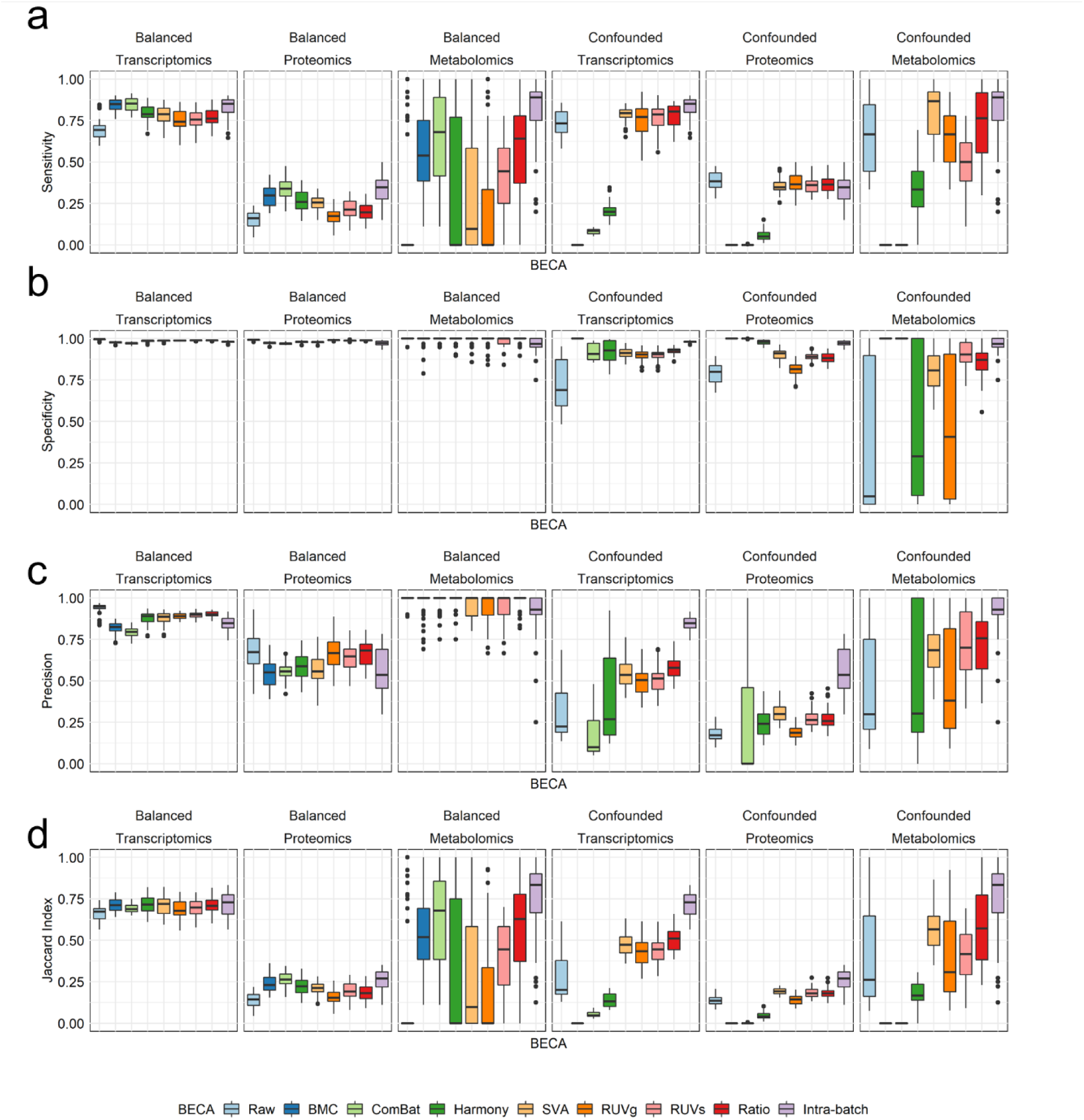
Evaluation of the performances of various BECAs based on the identified differentially expressed features. Sensitivity (**a**), specificity (**b**), precision (**c**), and Jaccard Index (**d**) of the DEFs.

**Extended Data Fig. 3.**
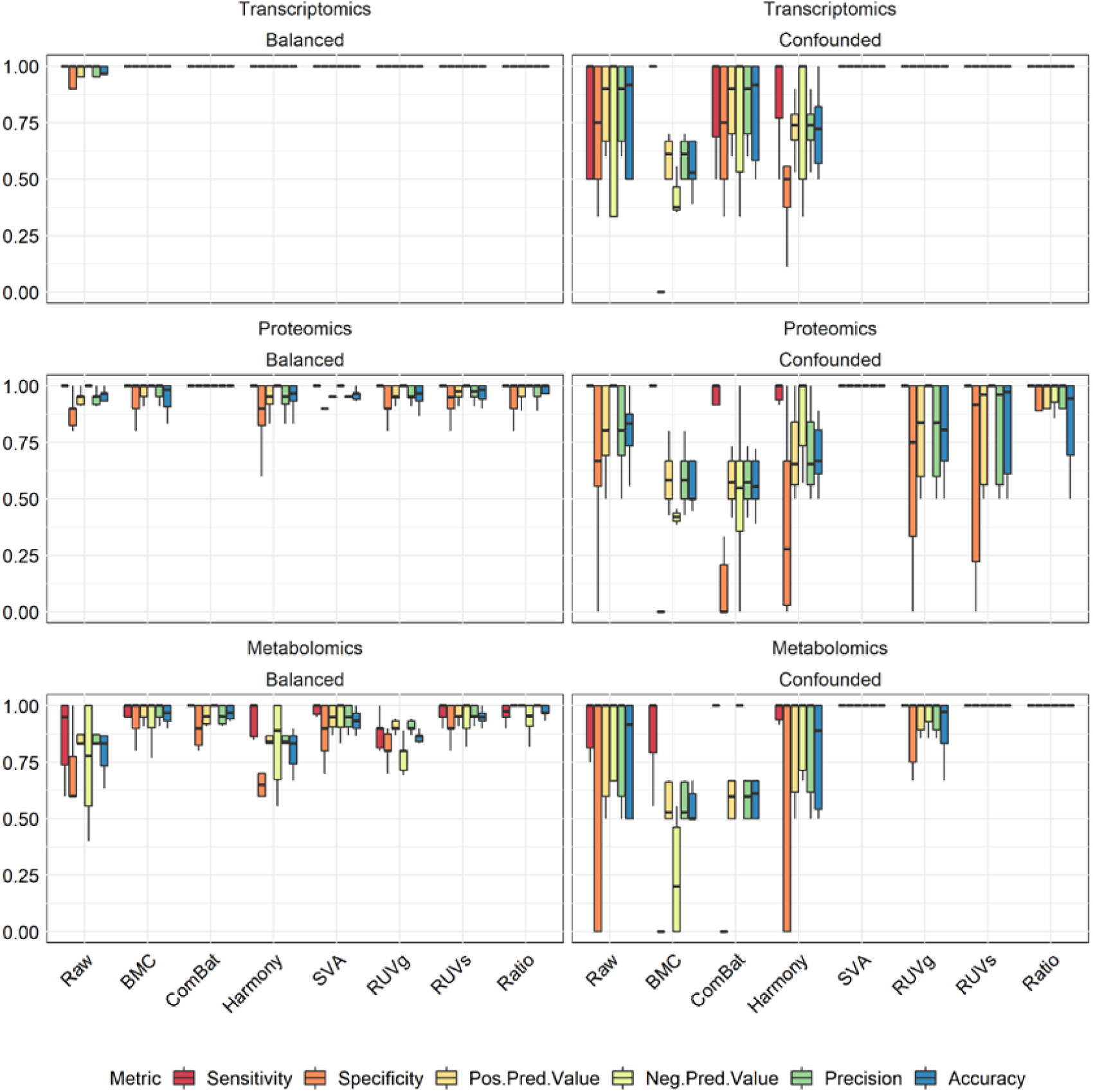
Evaluation of the performances of various BECAs based on model prediction results.

## Supplementary Information

**Supplementary Table 1. Description of datasets used in this study**

## Notes

### Competing Interest Statement

The authors have declared no competing interest.

